# Acute perilesional excitability explains long-term motor recovery after stroke

**DOI:** 10.64898/2026.02.27.708269

**Authors:** Julian Schulte, Gustavo Patow, Yonatan Sanz Perl, Jakub Vohryzek, Maurizio Corbetta, Gustavo Deco

## Abstract

Stroke is a leading cause of disability, and changes in neuronal excitability in perilesional regions may critically influence patients’ recovery. While reduced excitability is often associated with impaired recovery, increased excitability following injury may support reparative mechanisms. Although animal studies have shown how synaptic transmission adapts after stroke, a mechanistic understanding of how these excitability changes in the perilesional area relate to recovery in acute patients remains unclear. Here, we apply patient-specific (*N* = 96) computational whole-brain models to infer regional excitability, focusing on both perilesional and non-perilesional sites. Specifically, we estimate the firing sensitivity of local excitatory neuronal populations. Our findings indicate large inter-subject variability, with patients displaying both relative perilesional hypo- and hyper-excitability. Notably, perilesional excitability emerges as a robust predictor (*P* = 0.002) of motor recovery one year after the stroke, but not of acute post-stroke motor impairment, emphasizing its significance in specifically shaping long-term recovery. This synaptic modulation of excitability exhibits a strong correlation with gamma-aminobutyric acid A (GABA-A) receptor density distributions before the stroke, providing a potential biological substrate. These findings highlight the subject-specific nature of perilesional excitability, positioning it as a compelling target for personalized interventions to optimize post-stroke motor recovery.

## Introduction

Stroke remains a leading cause of disability and mortality worldwide, causing persistent impairments that profoundly affect patients and caregivers.^1^ While neuronal loss within the stroke core is irreversible, surrounding perilesional areas undergo functional and structural reorganization^2^, thus making these areas a potentially critical target for therapeutic interventions. Specifically, alterations in neuronal excitability are thought to be central to both impaired and successful recovery. Lesion studies in animals indicate that, following initial excitotoxicity, two opposing processes can occur simultaneously: a reduction in excitability coupled with increased growth-inhibiting factors that impede recovery; and an elevation in synaptic excitability, accompanied by growth-promoting factors that support functional restoration.^3^

The beneficial increase in excitability may arise from various synaptic alterations, such as reduced inhibition via downregulation of gamma-aminobutyric acid A (GABA-A) receptors^4–7^ or the suppression of inhibitory interneurons.^4,8^ Conversely, a direct increase of excitation can be the result of an upregulation of excitatory N-methyl-D-aspartate (NMDA) receptors.^9^ Consistent with the proposed benefits of enhanced excitation, pharmacological upregulation of (excitatory) α-amino-3-hydroxy-5-methyl-4-isoxazolepropionic acid (AMPA) receptor signalling has been shown to improve motor recovery^10^, as well as the attenuation of excessive extra-synaptic GABAergic tonic inhibition.^11,12^ Human non-invasive stimulation studies have mirrored these findings, demonstrating that increased excitation could be beneficial for recovery. Stroke patients have demonstrated both diminished ipsilesional motor cortex excitability but also generally reduced intracortical inhibition.^13^ Interestingly, this disinhibition^14^, alongside motor-evoked potentials^15,16^, have been shown to positively predict subsequent motor recovery. Furthermore, non-invasively stimulating neuronal excitability at the ipsilesional site has shown benefits for motor recovery.^17,18^

Although these studies underscore the critical role of neuronal excitability, little is known about the local underlying mechanisms in humans after stroke, and importantly, how they could relate to long-term recovery. Generally, magnetic resonance imaging (MRI) studies have primarily focused on structural damage topography^19,20^ and functional connectivity patterns^20,21^ to explain acute impairments. Regarding long-term recovery, relevant factors have included, amongst others, demographic and socioeconomic variables^22,23^, renormalization of functional connectivity^24^, structural lesion topography, and the integrity of the corticospinal tract.^23,25–27^ Beyond purely descriptive imaging, computational whole-brain modelling has proven valuable for uncovering the mechanisms driving functional connectivity alterations after stroke.^28–32^ Interestingly, in-silico lesion studies have shown that homeostatic adjustment of inhibitory synaptic weights can help restore functional connectivity features.^31,32^ Applying this modelling approach to actual chronic stroke patients, Falcon et al.^28^ demonstrated that a generally reduced excitatory drive onto inhibitory populations predicts better motor recovery, linking synaptic weight changes to therapeutic outcomes. While this previous work has underscored the role of excitability changes in stroke, it remains unknown how these alterations mechanistically unfold in the acute phase, particularly within perilesional cortex, an area critically involved in functional reorganization.

To address this critical gap, we employed personalised biophysical whole-brain models to estimate patient-specific perilesional excitability and its relationship to recovery trajectories. We simultaneously fitted the whole-brain models on patient-specific structural and functional features and investigated how the excitability i.e., the firing sensitivity of the excitatory neuronal populations changes. Our findings reveal that perilesional excitability is a clear predictor of motor outcomes at one year after the lesion, while it shows no relationship to acute motor impairments, highlighting its involvement in facilitating recovery. Furthermore, we found that perilesional excitability differs from non-perilesional excitability and exhibits substantial inter-patient variability, which remains stable over the course of one year. Additional analyses revealed that heightened perilesional excitability is associated with reduced pre-lesion GABA-A receptor densities, rather than structural characteristics of the lesion or overall structural damage, suggesting that it is mainly driven by local synaptic characteristics. Finally, we demonstrate that perturbing in-silico perilesional excitability at the acute stage approximates individual brain dynamics observed one-year post-stroke, underscoring its promise as a target for both pharmacological and non-invasive stimulation approaches.

## Materials and methods

### Subjects

The data for this study were obtained from the Washington University Stroke Cohort ^19^, a longitudinal dataset. We focused on measurements acquired at two time points: two weeks (T1) and one year (T3) post-stroke. The cohort predominantly comprised patients with ischemic strokes (83%), with the remaining 17% having suffered hemorrhagic strokes. For further details regarding data collection protocols, refer to the work from Corbetta et al..^19^ Patient data were obtained through the stroke service at Barnes-Jewish Hospital in collaboration with the Washington University Cognitive Rehabilitation Research Group, with a focus on patients experiencing a first-time, single-lesion stroke. This study included a total of *N = 96* stroke patients. For the calculation of the healthy effective connectivity map, we included 27 age-matched healthy individuals. Patients and controls gave informed consent, adhering to procedures approved by the Washington University Institutional Review Board. Comprehensive neuroimaging data, including both structural and functional measures, were acquired alongside detailed neuropsychological assessments.

### Neuropsychological and behavioural data

For each time-point, participants completed a battery of neuropsychological tests. A total of 44 measures were collected, spanning four key functional domains: motor control, language, attention, and memory function.^19^ For each functional domain, individual test data were subjected to dimensionality reduction using principal component analysis resulting in summary scores for: MotorR (right body side) and MotorL (left body side), language, MemoryS (composite spatial memory score), and MemoryV (composite verbal memory score), AttentionVF (visuospatial field bias), average attention performance (summarized performance score on the battery’s attention task), and AttentionValDis (the capacity to shift attention toward previously unattended stimuli). Motor tasks comprised both upper and lower body functions. Patients’ behavioural scores were normalized with z-scores relative to the control group, allowing for the identification of behavioural impairments. Overall composite scores were calculated by averaging the summary scores within each domain. Motor composite scores were derived from the combined MotorR and MotorL scores, while cognitive composite scores were obtained by averaging the individual summary scores from each cognitive domain.

### Neuroimaging acquisition and pre-processing

The MRI data were collected at the Washington University School of Medicine with a 12-channel head coil Siemens 3T Tim-Trio scanner. The detailed protocol for every study participant comprised of sagittal T1-weighted MP-RAGE (TR = 1950 ms; TE = 2.26 ms, flip angle = 90°; voxel dimensions = 1.0 × 1.0 × 1.0 mm), and a gradient echo EPI (TR = 2000 ms; TE = 2 ms; 32 contiguous slices; 4 × 4 mm in-plane resolution). Participants were instructed to maintain focus on a small white fixation cross displayed against a black background on a screen positioned at the rear of the magnet bore. 896 time points for each participants were sampled (~30 minutes) in between six and eight resting-state scans.^33^

The resting-state (f)MRI data were pre-processed following the procedure outlined below: (1) regression of head motion parameters, as well as signals from the ventricles, cerebrospinal fluid (CSF), and white matter; (2) a temporal filtering of the global signal maintaining frequencies between 0.008 – 0.08 Hz; (3) together with exclusion of frames with substantial head movements (FD = 0.5 mm). The filtered time-series was projected on 200 cortical and 34 subcortical & cerebellar regions of interest (ROIs). Cortical brain regions were parcelled according to the atlas of Schaefer et al. ^34^ and subcortical & cerebellar brain regions according to the automated anatomical labelling (AAL) atlas.^35^ For regions in the brainstem, the Harvard–Oxford Subcortical atlas (https://fsl.fmrib.ox.ac.uk/fsl/fslwiki/Atlases) was applied. As a last step, the BOLD signal was de-trended.

The structural connectome atlas was derived with diffusion MRI streamline tractography built upon the high-angular Human Connectome Project (HCP) diffusion MRI data sampled from 842 healthy participants.^36^ This atlas ^33,37^ was constructed in the MNI template space and is based on high-spatial and angular resolution diffusion imaging techniques and employs q-space diffeomorphic reconstruction.^38^ For all 842 subjects, the spin distribution functions were averaged to evaluate the diffusion patterns of the normal population-level. Whole-brain deterministic tractography was performed based on the population-averaged dataset, and multiple turning angle thresholds were applied to derive the 500000 population-level streamline trajectories. Following this procedure, the healthy controls group-averaged structural connectivity matrix *C* was constructed, indicating the anatomical connectivity weights between all brain regions.

### Stroke Lesions & disconnection masks

The manual segmentation of the structural MRI images was performed with the Analyze imaging software system^39^, and automatically categorized with an unsupervised K-means clustering approach according to their overlap with three masks (white matter, grey matter, and subcortical regions; see Corbetta et al.^19^). Perilesional brain regions were defined as those within 1 cm (Euclidean distance) surrounding the lesion, following previous research.^30,40^ For the acute stage (T1), we analysed the time series of 96 subjects. In six patients, the perilesional areas within 1 cm of the stroke core could not be clearly identified, and they were excluded from the relevant analysis. At the one-year follow-up, time-series data were available for 49 patients, of whom 45 had discernible perilesional areas. Across both time points, motor scores were obtained for 50 subjects, and 47 of these had identifiable perilesional regions. Overall, 72 participants had cortical perilesional areas (T1); because receptor-density data were available only for cortical ROIs, these individuals were included in the receptor-density–map analyses.

To evaluate the extent of structural connectivity damage, the Lesion Quantification Toolkit^37^ was applied, which estimates a collection of atlas-derived lesion measures including grey matter damage and white matter disconnections. The measures are inferred from population scale atlases of white matter connection trajectories and grey matter parcel boundaries.^36^ With the Lesion Quantification Toolkit, the structural disconnection masks (SDC) were estimated for each patient, indicating the percentage of intact structural connection between any two connected brain regions.

### Receptor density markers

For GABA-A and NMDA receptor density markers, we utilised the publicly available dataset by Hansen et al.^41^ where volumetric PET images were aligned to the MNI-ICBM 152 nonlinear 2009 template (version c, asymmetric), averaged across participants within each study, and subsequently parcellated. The individual GABA-A^42^ and NMDA receptor density markers^41,43–45^ were extracted for perilesional cortical brain regions, and subsequently the mean of them per subjects was calculated.

### Generative effective connectivity

Generative effective connectivity (GEC) is a model-based estimation of the directional influence of one brain region over another. It has been shown that enriching the GEC with the patient-specific SDC masks significantly contributed to the recovery of functional connectivity information in stroke patients.^29^ To leverage this information in our whole-brain models, we generated an average GEC from healthy controls (*N* = 27), following the methodology outlined by Deco et al..^46^ Subsequently, we performed an element-wise multiplication of it with the SDCs.

Specifically, to derive the GEC, the Hopf model was used to simulate BOLD activity, where each brain region is characterized by the normal form of a supercritical Hopf bifurcation and coupled via the empirical structural connectivity matrix *C*. The activity of a single brain region *j* is described as:

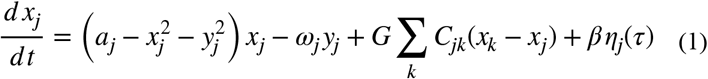

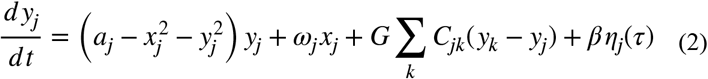

Where *x*_*j*_ describes the real (1) and *y*_*j*_ the imaginary part (2). Following previous literature, the bifurcation parameter *a* was set close to the Hopf bifurcation (*a* = ™ 0.01), and the intrinsic frequencies of the nodes denoted by *ω* were computed as the average peak frequency of the empirical narrowband BOLD signals associated with each brain region. The input from node *k* to *j* is scaled by the global coupling parameter *G* and *η* is a Gaussian noise vector with a standard deviation *β* = 0.04. The structural connectivity weights were scaled by 0.2, and optimization was performed by minimizing the difference of the phase of the model FC with respect to the phase of the empirical healthy FC (at *G* = 0.75).

After optimization of the whole-brain Hopf simulation, the GEC was computed by iteratively refining the initial structural connectivity weights *C* via a greedy gradient descent approach by assessing the distances between the model-based and empirical grand-average phase coherence matrices. Following previous studies, this process was repeated until the fit reached a stable value.^47–49^Specifically, weights in the connectivity matrix *C* were updated such that:

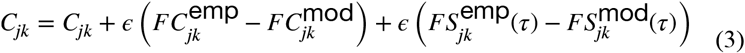

the difference between the grand-averaged empirical and model functional phase coherence matrices 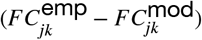 and the difference in time-shifted covariance matrices 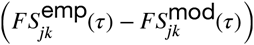 were scaled by *ϵ* = 0.001 and added to the previous connections. The covariance matrices were time shifted by the lag *τ* and normalized for each pair of nodes *i* and *j* by 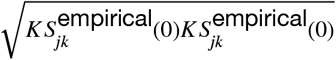. For further details, please refer to Idesis et al..^29^

### BEI dynamic mean field model

Regional brain activity is simulated using the Dynamic Mean Field model described by Deco et al..^50^ It is a biophysical whole-brain model describing spontaneous brain activity by the activation of local populations (here ROIs), which are connected via structural connections. Following the initial framework by Wong and Wang^51^, the model reduces the dynamics of large sets of interconnected excitatory and inhibitory integrate-and-fire spiking neurons into a system of equations comprehensively describing the activity of coupled excitatory (E) and inhibitory (I) populations. The activity in a single brain region is modelled by one excitatory and one inhibitory population, whose behaviour is shaped by their own recurrent activity and their interactions with each other. Whereas the excitatory synaptic currents are determined by *NMD* receptors, the inhibitory currents are determined by *GABA* receptors. Long-range activity between brain regions is propagated from one excitatory population to another, based on the connectivity weights in *C*_*ij*_ (here the GEC masked with the patient-specific SDC) and scaled by a global coupling factor *G*. The activity of a single brain region is *i* governed by:

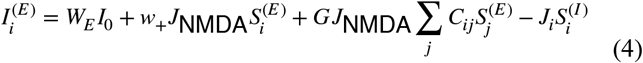

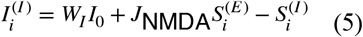

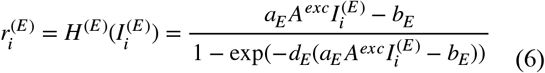

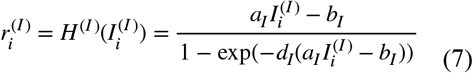

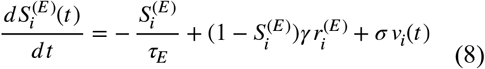

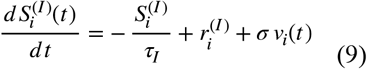

where the input current to the E/I population is denoted by 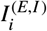, the firing rate by 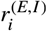, and the average synaptic gating by 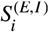. The general external input current *I*_0_ = 0.382(*n A*) is scaled by respectively *W*_*E*_ = 1 or *W*_*I*_ = 0.7. The excitatory synaptic coupling *J*_*NMDA*_ = 0.15(*n A*), and *w*_+_ = 1.4 is the local excitatory recurrence. *J*_*i*_ is the local feedback inhibitory synaptic coupling regulating excess excitation in the model and is estimated according to the computationally-efficient implementation of Herzog et al.^52^ by: 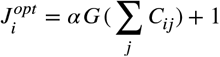, where *α* = 0.75.

*H* ^(*E*/*I*)^ are the neuronal input-output functions that transform the synaptic input currents to the population firing rates. To estimate the excitability of only the excitatory population, we scale the local excitatory current *I* ^(*E*)^ by *A*^*exc*^ in the first input-output function (equation 6). This parameter is later estimated and provides a general representation of the firing sensitivity of the excitatory population with regard to the synaptic input current, thus representing the population’s excitability. The other parameters in the input-output functions are static and listed in Table 1. The kinetic parameters are *τ*_*E*_ = *τ*_*NMDA*_ = 100*ms,τ*_*I*_ = *τ*_*GABA*_ = 10*ms*, and *γ* = 0.641/1000 (the factor 1000 for expressing everything in ms). Uncorrelated standard Gaussian noise *v*_*i*_(*t*) is added to the synaptic gating equations with an amplitude of *σ* = 0.01(*n A)*. The connectivity matrix *C* was scaled by 0.2 and for the numerical integration of the equations the Euler–Maruyama method was applied with an integration step of *dt* = 1.

**Table 1:** Biophysical parameters of the BEI model.

| Parameter | Value |
| --- | --- |
| $I_0$ | 0.382 (nA) |
| $W_E$ | 1 |
| $W_I$ | 0.7 |
| $J_{NMDA}$ | 0.15 (nA) |
| $w_+$ | 1.4 |
| $a_E$ | 310(nc<sup>-1</sup>) |
| $b_E$ | 125 (Hz) |
| $d_E$ | 0.16 (s) |
| $a_I$ | 615(nc<sup>-1</sup>) |
| $b_I$ | 177 (Hz) |
| $d_I$ | 0.087 (s) |
| $\tau_E = \tau_{NMDA}$ | 100 (ms) |
| $\tau_I = \tau_{GABA}$ | 10 (ms) |
| $\gamma$ | 0.641 / 1000 |
| $\sigma$ | 0.01 (nA) |

### Hemodynamic Model

The simulated excitatory firing rate 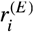 was transformed into BOLD signal by applying the hemodynamic Balloon–Windkessel model by Stephan et al..^53^ The vasodilator signal *s*_*i*_ for each brain region *i* is determined by the excitatory firing rate and subsequent auto-regulatory feedback. The resulting signal influences proportionally the differences in blood flow *f*_*i*_, leading to changes in blood volume *v*_*i*_ and deoxyhaemoglobin content *q*_*i*_. The mechanism is regulated by:

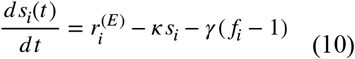

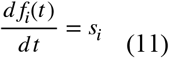

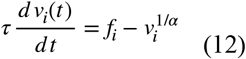

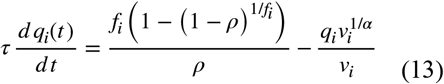

where *κ* is the rate constant of the vasodilatory signal decay, *γ* the rate constant for autoregulatory feedback by blood flow, *α* the resistance of the veins, *τ* a time constant, and *ρ* represents the resting oxygen extraction fraction. The blood volume *v*_*i*_ and the deoxyhaemoglobin content *q*_*i*_ subsequently determine the BOLD signal *B*_*i*_ :

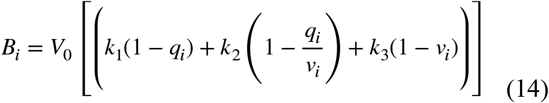

A more comprehensive description, along with all biophysical parameters used, is provided in Stephan et al.^53^ and Supplementary Table ST1. The simulated BOLD signals also underwent global signal regression, de-trending, and were band-pass filtered within the range of 0.008 to 0.08 Hz.

### Functional Connectivity and Functional Connectivity Dynamics

For the empirical and simulated BOLD signals, the functional connectivity (FC) was derived by calculating the Pearson correlation between each pair of brain regions. To capture dynamical features of the time-series we also calculated the functional connectivity dynamics (FCD). The FCD was calculated using a sliding-window approach (window size = 60 TR, step size = 12 TR), where windows were iteratively advanced, FC was computed within each window, and the resulting window-wise FC matrices were correlated with each other to obtain a final FCD matrix per patient.

### Model Optimization

The whole-brain model was optimized by estimating the global coupling parameter *G* and the excitability *A*^*exc*^. The excitability was estimated independently for perilesional brain regions (*Peri Exc*.) and non-perilesional brain regions (*Noperi Exc*.), constrained within the range *A*^*exc*^ ∈ [0.96,1.04] and *G* ∈ [0.1,12]. The three parameters were optimized by simultaneously fitting the simulated functional connectivity (Pearson correlation *ρ*) and functional connectivity dynamics (KS distance) to the empirical data. We implemented the particle swarm optimization (PSO) with a swarm size of 100 and a maximum iteration of 30 and performed it for the acute stage (T1) and follow-up (T3). The PSO aimed to minimize the model error, which was given by 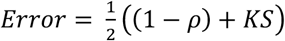. For the acute stage, results stabilized within 10–20 iterations, and the final parameter estimates yielded good model fits (Fig. S2). To increase the efficiency of the T3 optimization, we initialised the particles of the first iteration close to the optimal values obtained at T1. This led to initial low local minima while the PSO continued to search the entire range of the parameter space in the subsequent iterations. The PSO resulted likewise in a good model fit for the T3 optimisation (Fig. S4a, b). The Python implementation for the PSO can be found at: https://pythonhosted.org/pyswarm/.

### Statistical analysis

Acute (T1) and one-year (T3) motor as well as cognitive scores were predicted with multivariate linear regression analysis. Predictors included perilesional excitability, lesion etiology (i.e., ischemic or hemorrhagic), and expert localization (i.e., cortical, cortico-subcortical, subcortical, white matter only, brainstem, and other areas). Previous research highlighted the role of lesion topography in explaining acute motor and cognitive deficits and their degree of recovery.^19,23^ Therefore, we calculated the first five principal components of the link damage of all brain regions capturing 65% of the variance. Nodal link damage was initially calculated from subject-specific lesion masks by summing the link damages associated with each individual node. For the final model, p-values of individual predictors were corrected for false discovery rate (FDR) using the Benjamini–Hochberg procedure, and standardized coefficients were reported (for all standardized predictors, see Figure S3).

### Model Perturbations

For the perilesional perturbation, we decreased and increased the patient-specific perilesional excitability in the whole-brain model (T1) and evaluated the error of the resulting FC and FCD to the patient’s empirical ones after one year (T3). We evaluated perilesional excitability values in an interval of 3% above and below the optimal point at the acute stage (400 steps). The perturbation which minimised the error between the simulated and empirical FCD one year ahead was chosen as the optimal value. To establish a statistical benchmark, we selected the errors within a narrow range around the optimal points for T1 and for the approximated T3 (0.3% above and below; range of 40 steps; see also supplementary figure S5). For later comparison, we evaluated the means of these ranges within subjects.

## Results

The utilised stroke dataset comprised patients at the acute stage (T1; *N* = 96) and one year after the stroke lesion (T3; *N* = 49), comprising (f)MRI imaging data and behavioural outcomes (Fig. 1a). By implementing a biophysical whole-brain computational model^50^, we were able to estimate the excitability separately for perilesional and non-perilesional areas (Fig. 1b). Specifically, each brain region (ROI) is modelled as a circuit of interacting excitatory and inhibitory neural populations (Fig. 1c bottom; see Methods). To capture perilesional and non-perilesional excitability, we modelled the firing sensitivity of these respective areas. In detail, the excitability parameter governs the neural gain: it determines how easily an excitatory population increases its firing rate in response to a given synaptic input (Fig. 1c top; Methods). We estimated this as two uniform parameters per patient: one value shared by all perilesional regions and a separate value shared by all non-perilesional regions. Although specific to the excitatory population, this parameter effectively modulates the local excitation-to-inhibition (E/I) ratio (Supplementary Fig. S1). At the network level, these populations are globally connected via a healthy Effective Connectivity (EC) map, masked with subject-specific lesion information to account for structural disruptions (Fig. 1d left). To derive the excitability parameters from the patient data, we fitted them (together with a global coupling factor, G) to approximate the empirical Functional Connectivity (FC) and Functional Connectivity Dynamics (FCD; Fig. 1d), thereby accounting for both the overall co-activation of brain regions and the temporal state features of the BOLD signal.

**Figure 1.**
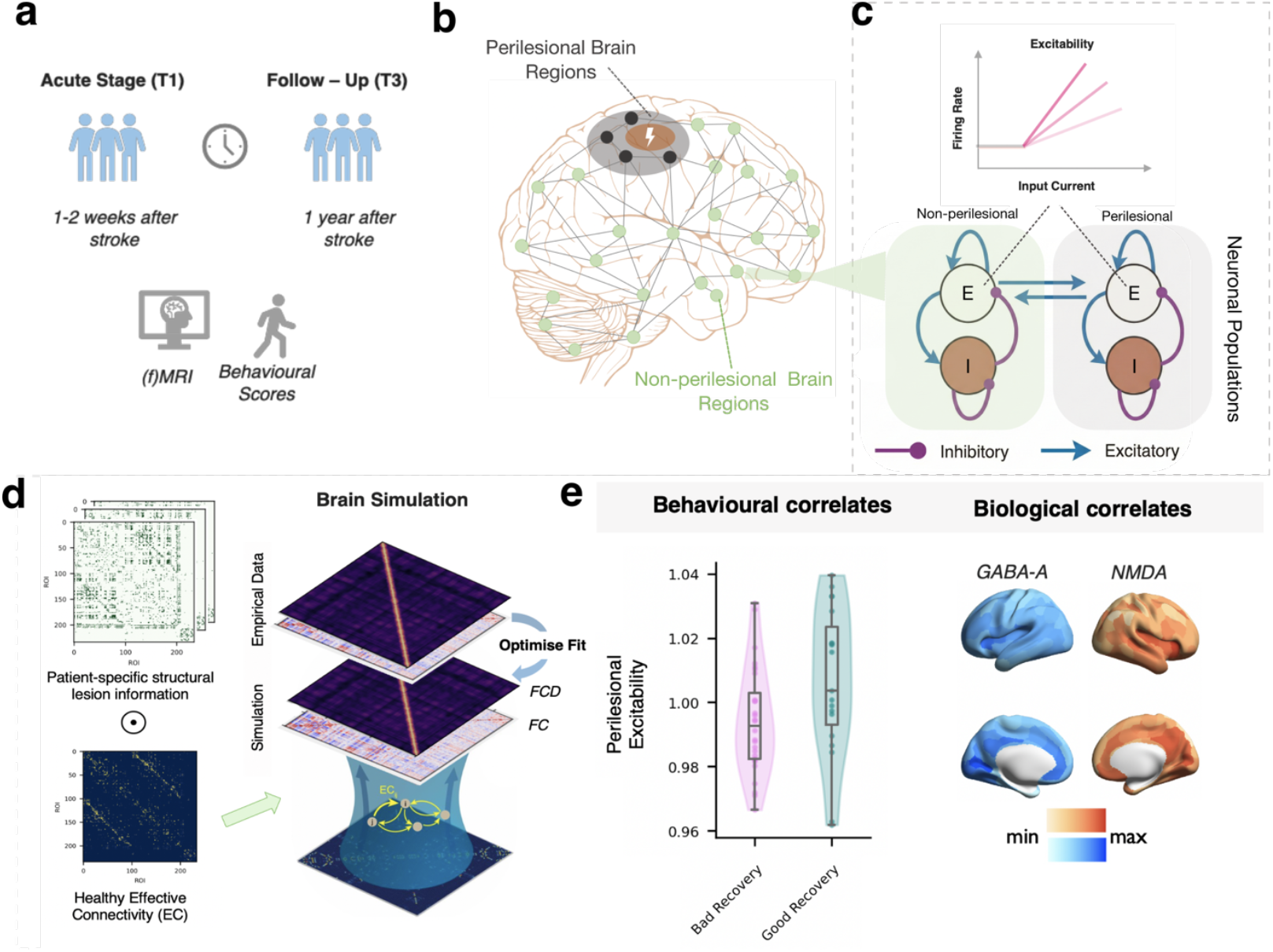
Whole-brain model pipeline. **a|**Imaging and behavioural data were analysed from stroke patients at the acute stage (T1; 1–2 weeks post-stroke) and one year post-stroke (T3). **b**|The overall goal of the patient-specific whole-brain simulations was to determine the neuronal excitability of perilesional (black dots) and non-perilesional brain regions (green dots). **c**|Local dynamics. Each brain region (ROI) is modelled as a circuit of interacting excitatory (E) and inhibitory (I) populations (bottom). To capture the excitability, we modelled the firing sensitivity of the perilesional areas (green) and non-perilesional areas (grey) as two independent parameters. Specifically, the excitability parameter governs the neural gain: it determines how easily an excitatory population increases its firing rate in response to a given synaptic input (top). **d** | Global dynamics and fitting. The connectivity between brain regions was defined by masking a healthy Effective Connectivity (EC) map with patient-specific lesion information (left). We then fitted the whole-brain simulation to each patient by optimizing the excitability parameters (simultaneously with the global coupling G). This optimization maximized the agreement between simulated and empirical Functional Connectivity (FC) and Functional Connectivity Dynamics (FCD), thereby capturing both the spatial co-activation of regions and the temporal state features of the BOLD signal. **e**|Subsequently, we examined whether individual estimates of perilesional excitability predicted behavioural outcomes and aligned with biological markers, including NMDA and GABA-A receptor density maps.

After fitting all three parameters, the model is able to capture the subject-specific features of the functional connectivity (Pearson correlation *M* = 0.36, *SD* = 0.068) as well as temporal dynamics of the BOLD signal (KS distance *M* = 0.12, *SD* = 0.062; Fig. S2). To elucidate the clinical role of perilesional excitability, we subsequently examined its relationship with behavioural scores and also explored potential biological correlates, including lesion characteristics and receptor density distributions in the perilesional cortex (Fig. 1e).

### Perilesional excitability predicts motor recovery one year after stroke

To determine how patient-specific perilesional excitability at the acute stage (T1) influences recovery, we next examined whether it predicts motor recovery at one year (T3). During this period, patients (*n* = 50) showed a significant improvement in motor scores from the acute stage (*T1*; *Mdn* = 0.302) to one-year post-lesion (*T3*; *Mdn* = 0.448), as indicated by the Wilcoxon signed-rank test, *W* = 210, *P* < .001 (Fig. 2a). As an initial analysis, stroke patients with good motor recovery (z-transformed recovery scores T3 – T1 > 0) exhibited significantly greater perilesional excitability (Peri Exc. T1; *n* = 19; *M* = 1.006, *SD* = 0.023) compared to those with poor recovery (*n* = 28; *M* = 0.994, *SD* = 0.016; *t*(45) = 2.13, *P* = 0.039]; Fig. 2b; Note: Sample sizes vary because the extend of motor scores and perilesional data were partly incomplete and differed between T1 and T3, see Methods - Stroke Lesions & disconnection masks).

**Figure 2.**
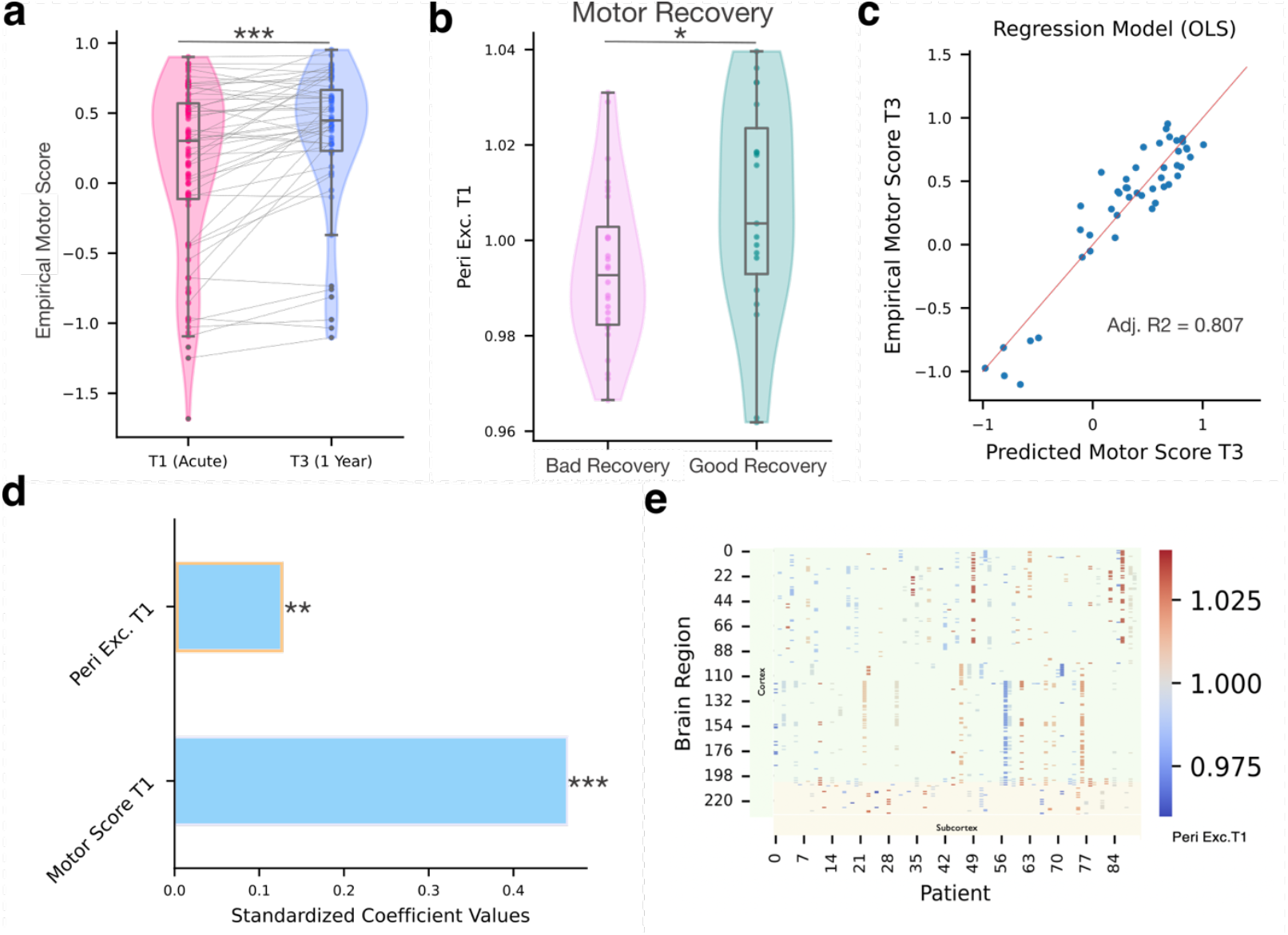
Acute (T1) perilesional excitability (Peri Exc.) predicts motor recovery after one year (T3) of the stroke lesion. **a**|Within-subject empirical motor scores improve after one year. **b**|Stroke patients with a good (z-transformed recovery scores T3 – T1 > 0) compared to a bad motor recovery show an increased Peri Exc. at T1. **c**|Accounting also for multiple factors, including initial motor impairment (T1), lesion type, expert localization, and structural connectivity damage, led to a high model fit of predicted motor scores at T3 (adj. R2 = 0.807). **d**| Among the significant predictors (not showing dummy variables; see Supplementary figure S3), Peri Exc. at T1 is a highly significant and clear predictor of motor scores at T3. **e**|Visualisation of final Peri Exc. at T1 for each patient coloured by subcortical (yellow) and cortical (green) brain regions. Statistical significance: *P < 0.05; ** P < 0.01; *** P < 0.001; n.s., not significant.

To further delineate the role of perilesional excitability in recovery, we performed a multivariate linear regression analysis, incorporating additional factors that could influence recovery trajectories (see Methods). The model explaining one-year motor scores (T3) was statistically significant (*F*(14, 32) = 14.73, *P* < 0.001, *adjusted R*^2^ = 0.807; Fig. 2c) and included as predictors the perilesional excitability, initial motor impairment (T1), lesion type (ischemic/ hemorrhagic), expert localization (e.g. cortical, cortico-subcortical, etc.), and five principal components accounting for structural connectivity damage per brain region (see supplementary figure S3 for complete model). Importantly, perilesional excitability was a highly significant positive predictor of motor recovery (standardized *β* = 0.126, *SE* = 0.038, *t*(32) = 3.298, *P* = 0.002, *95% CI* [0.048, 0.204]; Fig. 2d), and this association remained significant after correcting for False Discovery Rate using the Benjamini-Hochberg method (FDR-corrected *P* = 0.009), showing that higher excitability was associated with greater improvements in motor function.

We also performed multivariate linear regressions to predict acute motor and cognitive scores (T1) and one-year cognitive outcomes; however, perilesional excitability was not a significant predictor. Overall, this analysis revealed that perilesional excitability is a specific and clear predictor of motor outcomes at one-year post-stroke, suggesting a particular role in long-term motor restoration.

### Perilesional excitability is independent from non-perilesional excitability and shows high heterogeneity across patients

After evaluating the influence of perilesional excitability on functional recovery, we next sought to better understand how it relates to the excitability of other brain regions and the global coupling parameter. For the acute stage, there was no significant correlation between peri- and non-perilesional excitability (Spearman’s *r(88)* = −0.08, *P* = 0.44), highlighting distinct excitability profiles in regions adjacent to the stroke lesion compared to more distant areas (Fig. 3a). Similarly, the global coupling parameter was not correlated with the perilesional excitability (Fig. 3b; Spearman’s *r(88)* = 0.09, *P* = 0.38), but showed a strong negative correlation with non-perilesional excitability (Fig. 3c; Spearman’s *r(94)* = −0.76, *P* < 0.001). This inverse relationship suggests a trade-off between local excitation of non-perilesional brain regions and the global scaling of excitatory signalling between brain regions.

**Figure 3.**
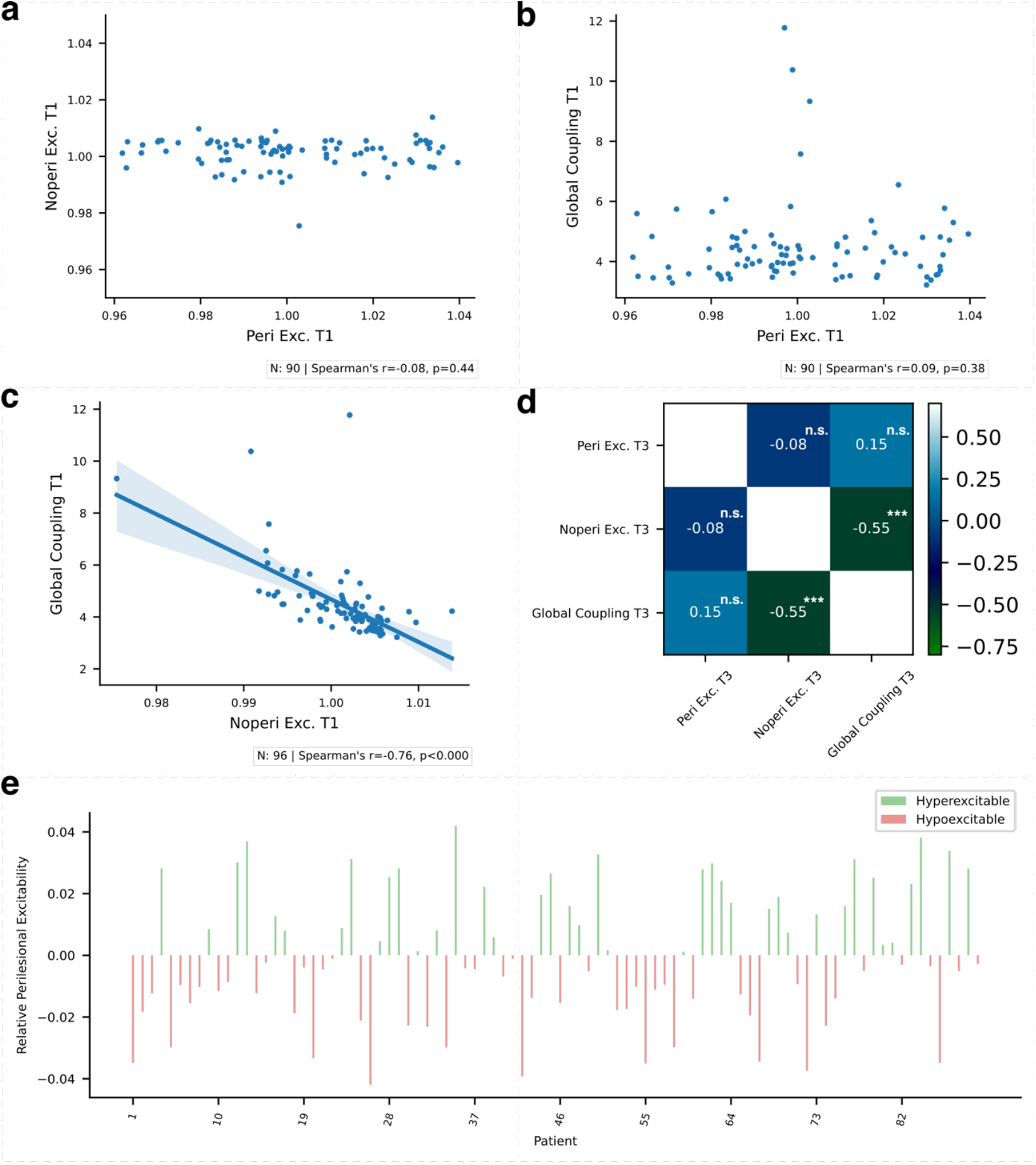
The excitability near the lesion is distinct from both the excitability in non-lesioned areas and the overall global coupling G. **a**|Perilesional excitability (Peri Exc.) is uncorrelated to non-perilesional excitability (Noperi Exc.) **b**|and also to the global coupling factor G. **c**|However, Noperi Exc. is negatively correlated with the global coupling factor G. **d**|Estimating Peri Exc., Noperi Exc., and G for patients one year after stroke (T3), as well only Noperi Exc. and G were negatively correlated. **e**|Patients widely differ in their relative perilesional excitability at T1 (i.e., the fraction of Peri Exc. to Noperi Exc. - centred at zero) showing different hypo- and hyperexcitability profiles.

To evaluate how these parameters change over time, we subsequently estimated them one year following the stroke lesion. Similar to the acute stage, only non-perilesional excitability and the global coupling parameter exhibited an inverse correlation (Fig. 3d). We next assessed the temporal stability of the parameters over the one-year period. Overall, the parameters exhibited high stability throughout this timeframe. Perilesional excitability showed a strong correlation between the acute stage and follow-up (Pearson’s *r(43)* = 0.92, *P* < 0.001; Supplementary Fig. 4c), followed by the global coupling (Spearman’s *r(47)* = 0.50, *P* < 0.001; Supplementary Fig. 4e) and non-perilesional excitability (Spearman’s *r(47)* = 0.40, *P* < 0.01; Supplementary Fig. 4d).

To elucidate whether the perilesional areas are relatively hyper- or hypo-excitable as compared to other brain regions, we calculated an excitability ratio for the acute stage by dividing perilesional excitability by that of non-perilesional regions. Interestingly, patients showed both hyper- and hypo-excitability of the perilesional areas (Figure 3e; excitability ratio was centred at zero). Overall, these findings suggest that excitatory neuronal populations in perilesional areas maintain stable excitability over time, exhibiting distinct profiles compared to non-perilesional regions, with subject-specific variability.

### Perilesional excitability is inversely related to local GABA-A receptor density but not lesion or connectivity features

While perilesional excitability serves as a predictor of long-term motor recovery, potential factors modulating excitability remain unclear. We first examined the relationship between perilesional excitability and the density of cortical *pre-lesion* inhibitory (GABA-A) and excitatory (NMDA) neurotransmitter receptors (Fig. 4a). Our analysis revealed a significant inverse correlation between GABA-A receptor densities and perilesional excitability (Spearman’s *r(70)* = −0.34, *P* = 0.003, *n* = 72; Fig. 4b). This relationship indicates that higher pre-lesion GABA-A receptor densities within the cortical perilesional areas are associated with reduced excitability of the excitatory neuronal populations. Contrary to our expectations of a positive relationship, we observed a similar yet insignificant trend of inverse correlation between NMDA receptor density and excitability (Spearman’s *r(70)* = −0.23, *P* = 0.051, *n* = 72; Fig. 4c).

**Figure 4.**
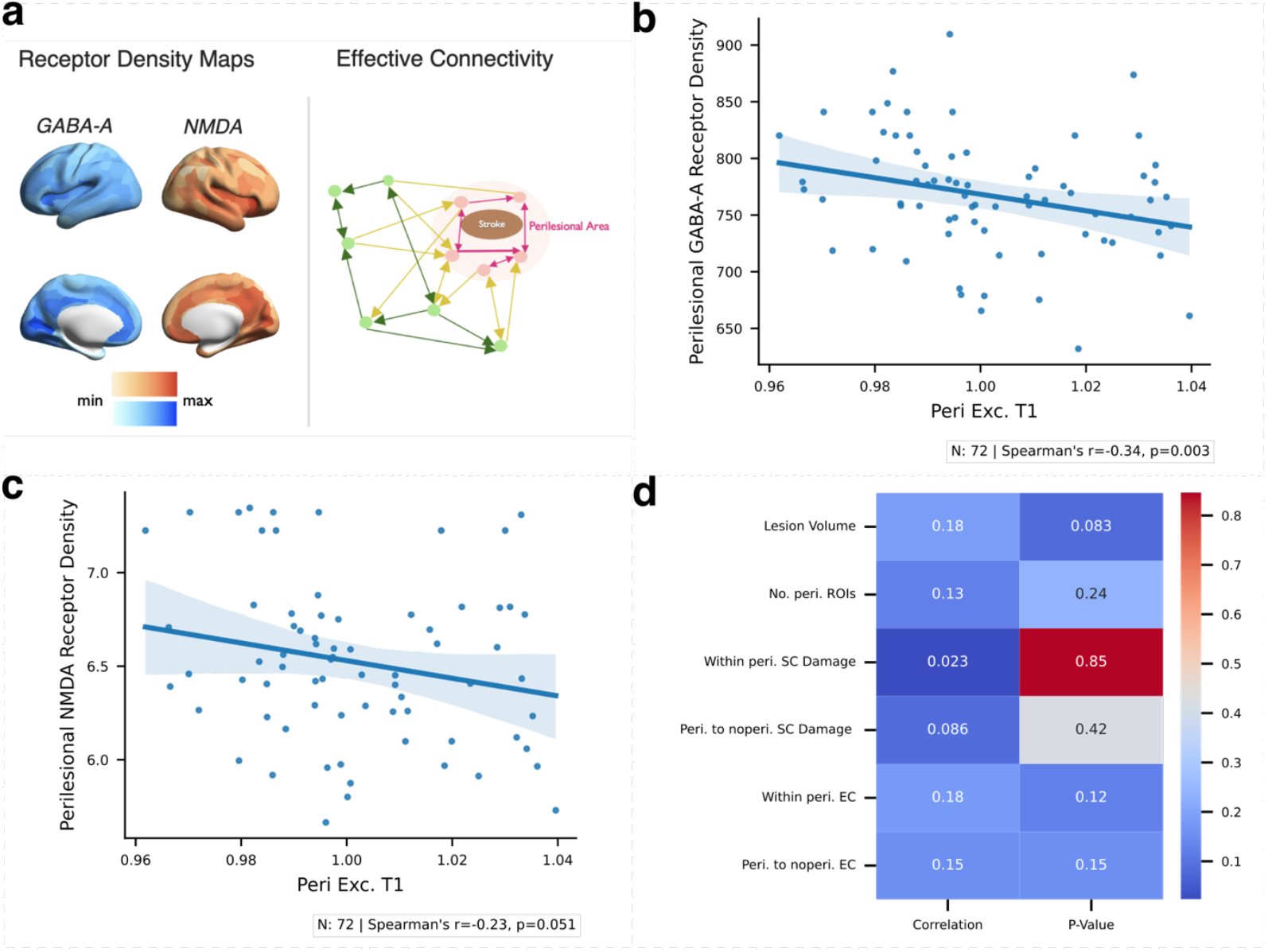
Heightened perilesional excitability is linked to reduced pre-lesion cortical GABA-A receptor densities but not to SC, EC, or general stroke lesion features. **a**|**Left** Visualization of cortical GABA-A (blue) and NMDA (orange) receptor density distributions. **Right** Effective connectivity (EC) weights between non-perilesional brain regions (green), non-perilesional to perilesional brain regions (yellow), and between perilesional brain regions (red). **b**|Perilesional excitability significantly inversely correlates with pre-lesion GABA-A receptor densities of cortical perilesional ROIs, **c** |and a tendency of this relationship is also observed for NMDA receptor densities. **d**|Lesion, SC, and EC features are not significantly correlated (Spearman’s r) with perilesional excitability.

We next investigated whether lesion extent or connectivity metrics might account for perilesional excitability. Specifically, we examined stroke lesion volume, the number of perilesional brain regions, structural connectivity (SC) disruption within perilesional areas or between perilesional and non-perilesional regions, and effective connectivity (EC) strength within and from perilesional to non-perilesional areas (Fig. 4d). Interestingly, none of these measures displayed a significant relationship, indicating that while perilesional excitability is associated with reduced pre-lesion GABA-A receptor densities, it is not primarily driven by local structural and effective properties or overall lesion extent.

### Directed perturbation of perilesional excitability can approximate functional connectivity and dynamics one-year post-lesion

Following a stroke, most patients exhibit some degree of motor recovery within one year (Fig. 2a). As perilesional excitability emerged as a key predictor of successful motor recovery, we investigated whether perturbing this excitability *in-silico* at the acute phase (T1) — by either increasing or decreasing it (see Methods) — could mimic patient’s joint functional connectivity (FC) and its dynamics (FCD) observed one-year post-lesion (T3; Fig. 5a). We also evaluated how these changes relate to the degree of motor recovery only considering recovering patients.

**Figure 5.**
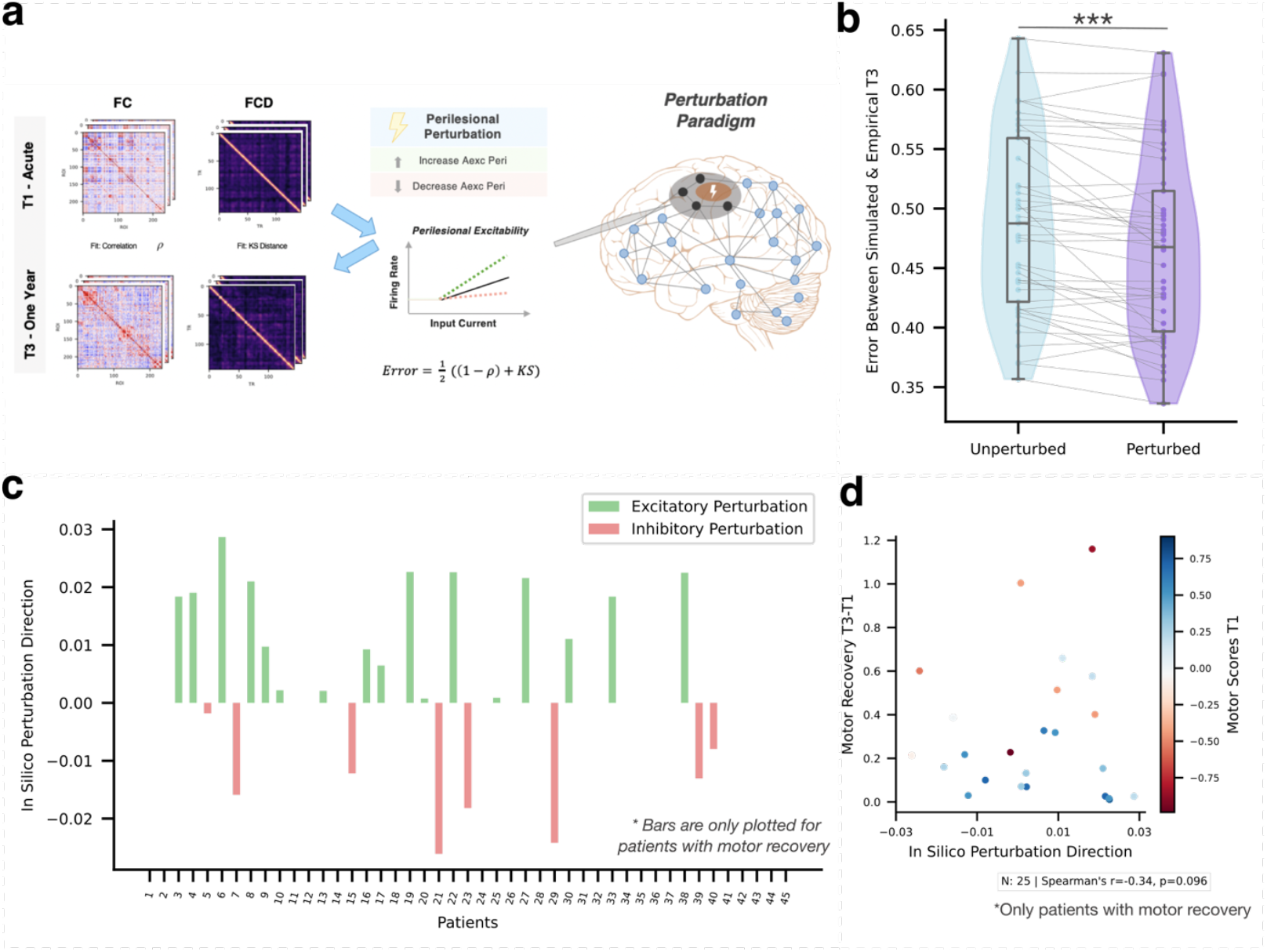
Patients’ joint functional connectivity and functional connectivity dynamics can be perturbed to resemble those observed one year later. **a**|We performed in-silico perturbations by in- and de-creasing the individually estimated perilesional excitability (Peri Exc.) for T1. For each perturbation, we calculated the resulting FC and FCD and compared their joint distance (i.e., error) to the patient’s empirical FC and FCDs after one year (T3). Whereas some patients required an increase (middle – green line) or a decrease (middle – red line; supplementary figure S5) of Peri Exc. to approximate the functional connectivity and dynamics of T3, some patients did not show clear differences. **b**|We considered a close interval around the initially estimated excitability (for T1) and for the excitability that minimizes the error for T3, and compared the mean values of these intervals. By finding an optimal value for Peri Exc., the errors for T3 can be significantly reduced. **c**|Bars are plotted only for patients exhibiting motor recovery greater than zero. Both excitatory (green) and inhibitory (red) optimal perturbations were observed in approximating their empirical FC and FCD at T3. **d**|The in-silico perturbation direction is uncorrelated to the degree of empirical motor recovery (for patients with at least minimal motor recovery).

Examining the perturbations for individual patients, we found that while some showed little change, others required either an in- or de-crease of perilesional excitability to approximate the joint FC and FCD after one year. For each patient, we identified the perturbation that best aligned with their empirical T3 data. As the error fluctuated around this optimal perturbation, we considered a narrow range of nearby perturbation values and their corresponding errors (see Methods & supplementary figure S5). To establish a reference point for comparison, we calculated the error associated with each patient’s initial excitability estimate for the acute stage (T1), including also a small surrounding interval. We then assessed whether the optimal perturbation brought the joint FC and FCD closer to the one-year follow-up state (T3). Specifically, we compared the mean error within the optimal perturbation interval (Fig. 5b, violet violin; *Mdn* = 0.468) to that of the interval around the initial T1 excitability (Fig. 5b, blue violin; *Mdn* = 0.488), within each subject. A Wilcoxon signed-rank test showed that these optimal perturbations significantly approximated the joint FC and FCD at T3 compared with T1 (*P* < 0.001, *z* = –4.261, *W* = 140.0).

Restricting our analysis to patients who showed motor recovery (i.e., Motor T3 – T1 > 0) revealed both excitatory and inhibitory optimal perturbation directions, defined as relative increases or decreases in excitability compared to T1 (Fig. 5c). However, perturbation direction did not correlate with the extent of motor recovery (Spearman’s *r*(23) = –0.34, *P* = 0.096; Fig. 5d). These findings suggest that shifting perilesional excitability, whether via up- or downregulation, can steer brain functional connectivity and dynamics toward patterns observed one year after lesion onset. However, both excitatory and inhibitory in-silico perturbations are associated with varying degrees of motor recovery across patients, underscoring the complex and nuanced relationship between functional brain state alterations, excitability, and motor recovery.

## Discussion

A mechanistic understanding of how perilesional excitability changes and consequently supports functional recovery has remained elusive in acute stroke patients. In this study, we have bridged this gap by computationally estimating subject-specific perilesional excitability using a biophysical whole-brain model that utilised individual functional and structural damage information. Our findings demonstrate substantial heterogeneity in perilesional excitability among stroke patients and establish this marker as a robust predictor of motor recovery one-year post-stroke. As a biological correlate, perilesional excitability displayed an inverse association with pre-lesion GABA-A receptor densities, yet remained unaffected by structural lesion features, underscoring the selective influence of receptor distributions on this marker. We further performed in-silico perturbations of patient-specific perilesional excitability and demonstrated that these changes alone could approximate the functional connectivity and temporal dynamics measured one year after stroke. This underscores the importance of localized excitability in long-term recovery and suggests that perilesional excitability could be a promising target for personalized therapies.

These findings provide further support in human patients aligned with previous research, showing that reduced neuronal excitability hinders functional recovery, while increased excitability promotes it.^3^ We demonstrated the clinical relevance of model-driven estimates of perilesional excitability, as they offer a mechanistic explanation and serve as a robust positive predictor of long-term motor recovery in acute stroke patients. These results align with a previous experimental non-invasive brain stimulation study showing that greater acute disinhibition of the ipsilesional motor cortex, likely driven by the GABAergic inhibitory system, is associated with better motor recovery three months after stroke.^14^ Interestingly, perilesional excitability did not predict initial motor impairments, underscoring its specific relevance for the recovery process. Therefore, this model-driven marker complements other predictors—such as the initial motor impairment, the location and topography of structural damage, the disinhibition of the ipsilesional motor cortex, and the integrity of white matter pathways, particularly the corticospinal tract^14,16,23,25–27^—enhancing the estimation of motor recovery trajectories. Regarding the recovery process for chronic stroke patients, previous research demonstrated that synaptic characteristics also play a crucial role in the long-term. In particular, Falcon et al.^28^ showed that the overall synaptic weight of excitatory over inhibitory populations was inversely related to motor performance after therapy.

Interestingly, computational studies have shown that specifically the synaptic scaling of inhibitory synapses in form of homeostatic plasticity can restore healthy functional connectivity^31,32^ but not dynamic functional connectivity features.^31^ In animal models several synaptic changes were observed that could contribute to a change of excitation: after induced stroke, an upregulation of NMDA receptors^9^ was observed but also a downregulation GABA-A receptors^4–7^, together with a lower expression of inhibitory interneurons.^4,8^

To elucidate the underlying synaptic mechanisms of the perilesional excitability, we examined its correlation with markers of GABA-*A* and NMDA receptor densities in the cortex of healthy individuals. We found a negative association with GABA-A receptor density distributions. While a higher presence of GABA-*A* receptors is intuitively associated with reduced excitability, it is notable that this relationship persisted even in the post-stroke lesioned state, as changes in GABA-A receptor expression are reported to take place after brain injury.^4–7,11,54^ Although our analysis could not account for post-lesion changes in receptor regulation, our findings suggest that the pre-lesion distribution of GABA-A receptors remains a significant determinant of perilesional excitability. In contrast, the pre-lesion distribution of NMDA receptors might *not* be decisive in determining the perilesional excitability. While we hypothesized a positive correlation between NMDA receptor distribution and perilesional excitability, our analysis revealed a statistically insignificant indication of an inverse relationship. The actual change of receptor regulation might be more relevant, as a previous study found a widespread, partly temporally delayed, upregulation of NMDA receptors.^6^

We also tested whether lesion characteristics or structural and effective connectivity relate to perilesional excitability, but found no significant correlations. Likewise, Falcon et al.^28^ reported that the size and location of the stroke in chronic patients did not correlate with model-estimated synaptic weights. While there might be other, unexamined aspects of structural damage or lesion characteristics, which may play a more prominent role, these findings suggest that receptor distributions and probably changes of those are more relevant determinants.

As motor function typically improves over time, we demonstrated that perilesional excitability can be modulated to approximate the joint FC and FCD observed one-year post-lesion. Among patients with at least some degree of motor recovery, responses varied, with some showing a better approximation from inhibitory and others from excitatory perturbations. Interestingly, no differences in motor improvement were observed related to whether the stimulation was excitatory or inhibitory, hinting that patients might benefit from different perturbation paradigms. It remains to be investigated whether modulating FC and FCD towards the state observed one-year post-lesion is beneficial or if the acute excitability levels are already optimal and should be preserved. Nevertheless, excitability-promoting stimulation paradigms have been shown to likely support motor recovery.^17,18,55^ In this line, our framework could also potentially enable patient stratification by identifying individuals with severe hypo-excitability who may benefit from excitatory stimulation, while reducing the risk of overstimulation in those with elevated perilesional excitability. Pinpointing the optimal perturbation target and validating it in clinical studies in a systematic way are the essential next steps for translating these model-informed results into clinically actionable outcomes.^56,57^

While local perilesional excitability strongly predicted the extent of motor recovery over one year, it showed no predictive value for acute cognitive impairment or long-term recovery. One possible explanation is that the degree of recovery for cognitive processes relies more on distributed than local compensation mechanisms. Siegel et al.^20^ found that visual and motor impairments were best explained by the corresponding lesion location. However, attention and memory deficits were best explained by functional connectivity and in particular by functional connections that linked various brain systems.^20^ Congruently, the re-emergence of network modularity has been associated with improvements in language, memory, and attention, but not with visual or motor deficits^58^, highlighting the strong network dependence on the recovery of these cognitive domains.

A limitation of this study is that the dataset primarily includes ischemic stroke patients, and validation in hemorrhagic stroke is needed to generalize these findings across all stroke types. Also, we estimated perilesional excitability in patients in the acute stage. Although this is a relatively short window, excitability may fluctuate during this period, particularly in the first few days. Thus, distinguishing between the early detrimental effects of excitotoxicity and the potential positive effects of later increased excitability is challenging, which may have introduced variability into the dataset. Next, in our analysis, we assessed motor function using a single, albeit comprehensive motor score. Future studies could further explore which specific motor functions may, or may not, benefit from increased perilesional excitability. Lastly, we focused on the excitability of the excitatory neuronal population to delineate the problem. Although this excitability marker directly influences the excitatory-inhibitory ratio (see Supplementary Figure S1), future studies should also investigate the perilesional excitability of the inhibitory neuronal populations in acute stroke patients.

Despite these limitations, this study provides new insights into patient-specific neural mechanisms of post-stroke recovery, showing that acute perilesional excitability is associated with long-term motor outcomes. By bridging computational neuroscience with clinical research, this work contributes to a mechanistic understanding of perilesional excitability in relation to motor recovery and opens promising avenues for the development of personalized therapeutic strategies.

## Data availability

Data can be made available upon request.

## Acknowledgments

We would like to thank Martí Monge-Asensio and Gorka Zamora-López for their helpful discussions and for providing useful suggestions regarding parts of the analysis. The open source software FieldTrip^59^ was used for the visualization of cortical receptor density maps.

