## Supplementary Materials for "Acute perilesional excitability explains long-term motor recovery after stroke"

| Parameter | Value |
|---|---|
| $\kappa$ | 1 / 0.65 |
| $\gamma$ | 1 / 0.41 |
| $\tau$ | 0.98 |
| $\alpha$ | 0.32 |
| $k_1$ | $4.3\mathscr{V}_0 E_0 TE$ |
| $k_1$ | $r_0 E_0 TE$ |
| $k_1$ | 1 |
| $\mathscr{V}_0$ | 40.3 |
| $E_0$ | 0.4 |
| $TE$ | 0.04 |
| $r_0$ | 25 |

**ST1 | Biophysical parameters of the Hemodynamic Model**

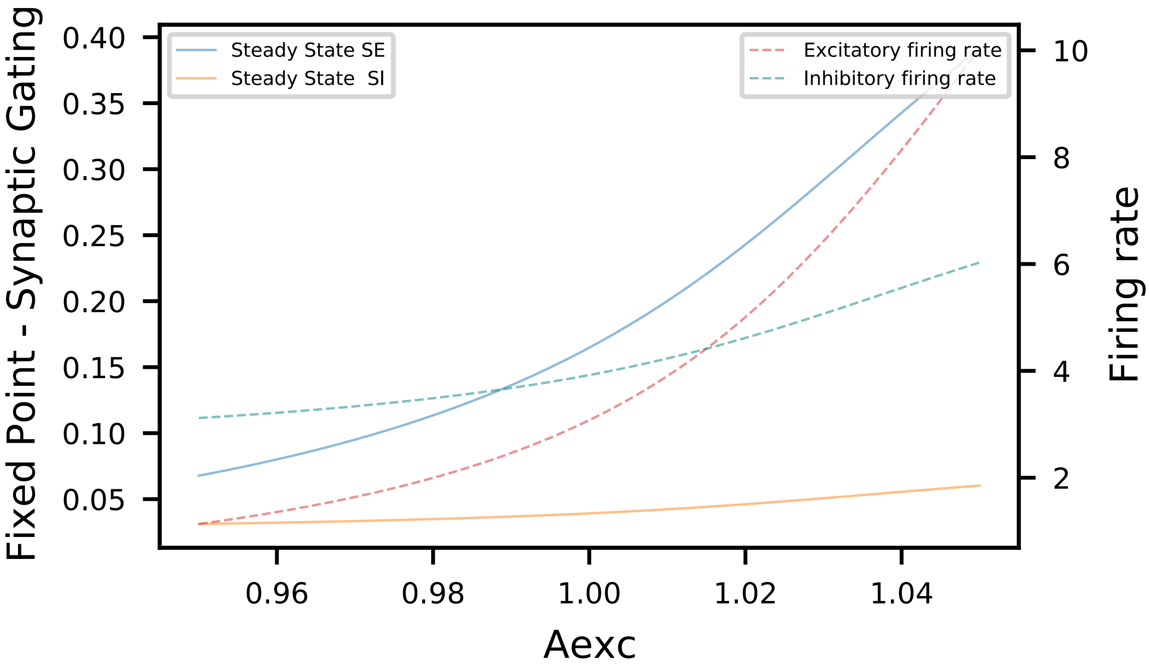


***Figure S1 -*** *Single Node Steady-State Dynamics of Synaptic Gating Variables | As excitatory excitability* $A^{exc}$ *increases, the steady-state values of the excitatory population's synaptic gating variable* $S^{(E)}$ *also rises, though with greater magnitude as compared to the inhibitory population* $S^{(I)}$*. Initially, the node exhibits inhibitory dominance; however, with increased excitability, the node shifts toward an excitatory firing rate dominance.*

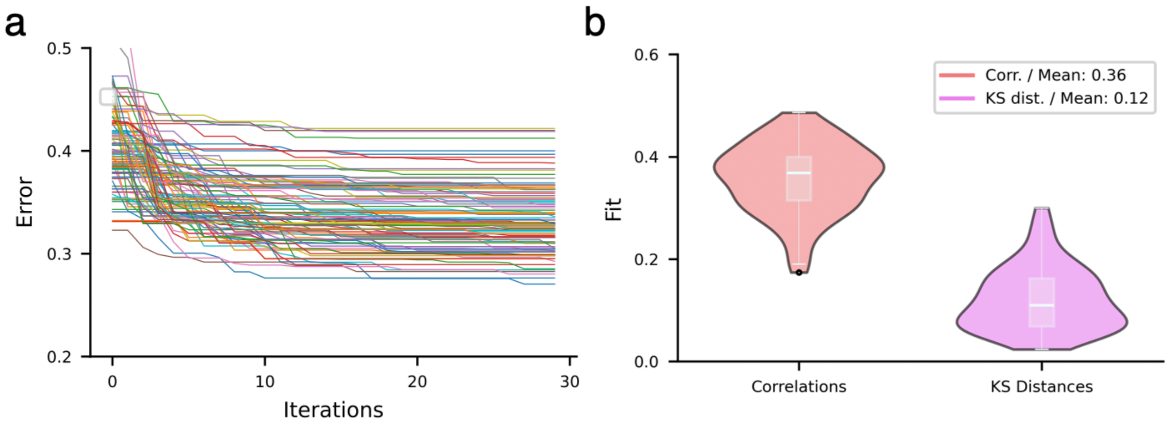


***Figure S2*** *– Particle Swarm Optimization (PSO) results – T1.* ***a****| The PSO minimised the errors for the patient-specific simulations over 30 iterations. Most results stabilized between 10 and 20 iterations.* ***b****| The final parameter fits of the PSO optimization yielded good correlations for the FCs and low KS distances for the FCDs between simulated and empirical data, indicating appropriate model fit.*

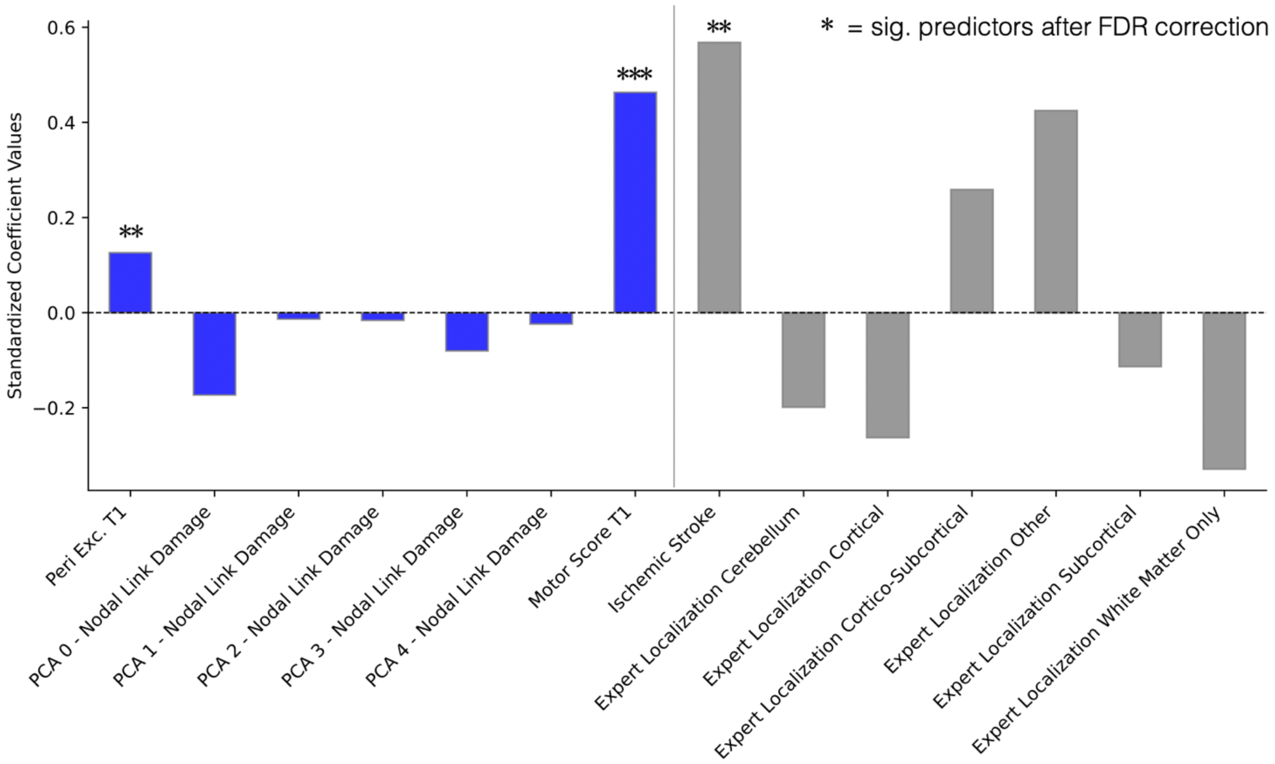


***Figure S3*** *– Standardized coefficients of the regression model predicting motor scores one-year post stroke (T3). Blue bars represent the standardized coefficients of continuous independent variables, while grey bars correspond to the estimated coefficients of categorical (dummy) variables. The reference categories are ischemic stroke for stroke type and brainstem for expert localization. Asterisks denote predictors that remain statistically significant following false discovery rate (FDR) correction.*

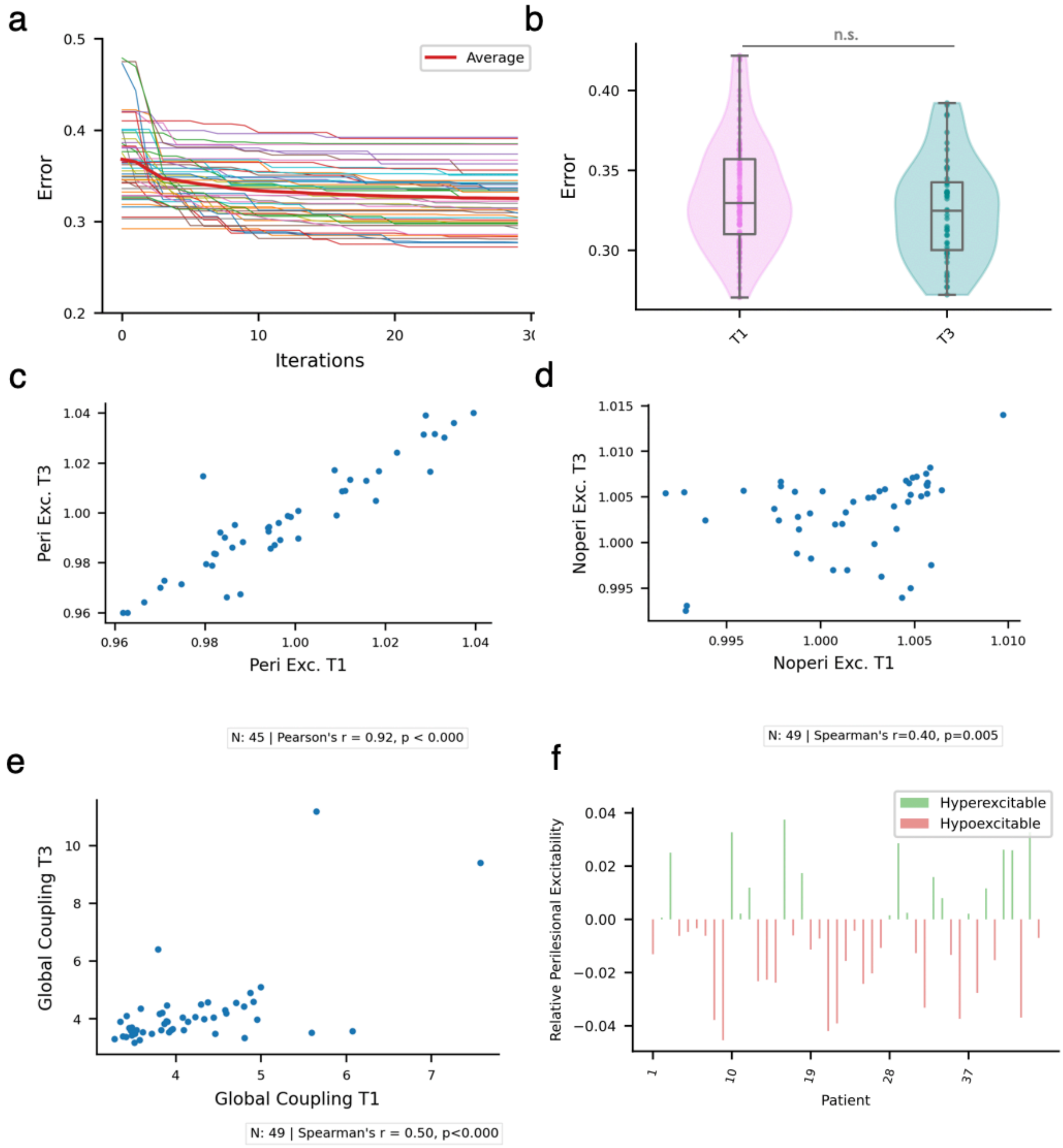


***Figure S4 Fitting and parameter results for one-year after the stroke lesion (T3) a***| *PSO optimization error for all 30 iterations.* ***b*** *| The final model error of T3 per subject is not significantly different from that of T1.* ***c*** *| Perilesional excitability of T1 and T3 are highly correlated similarly as* **d** *| the non-perilesional excitability and* ***e*** *| the global coupling factor G.* ***f*** *| Relative perilesional excitability (perilesional divided by non-perilesional excitability and centred at 0) of the patients at T3.*

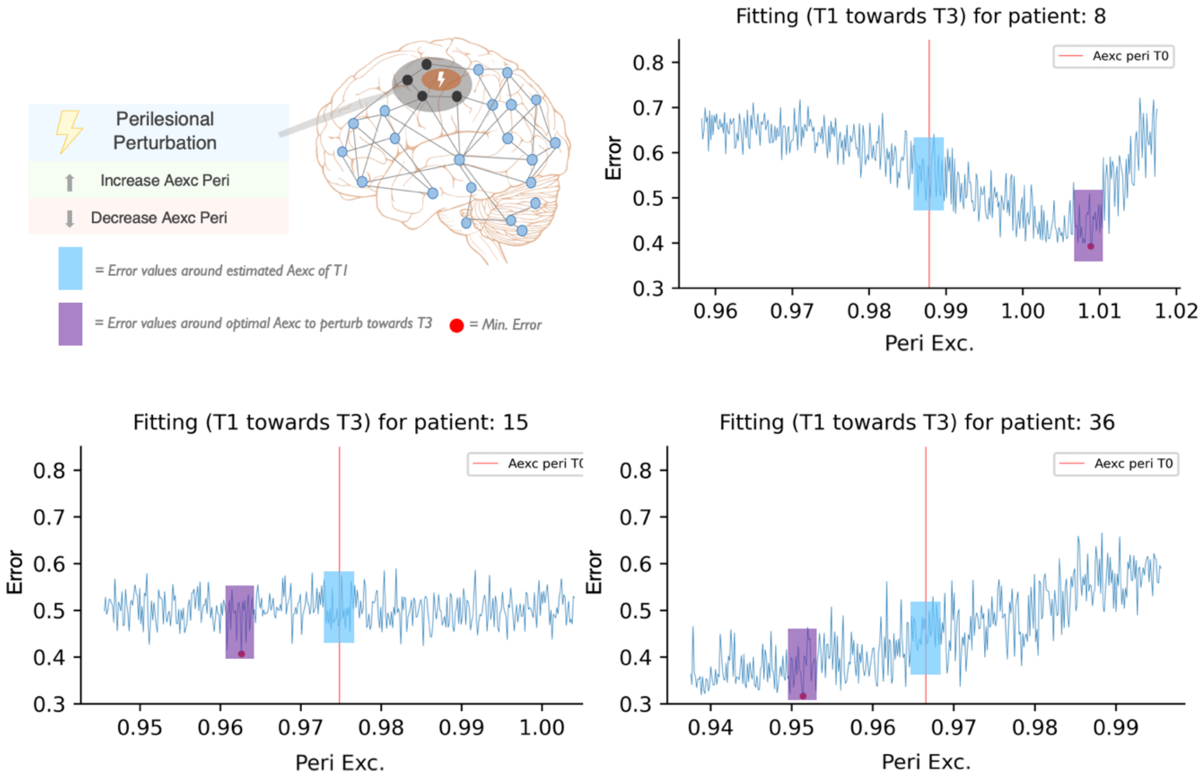


***Figure S5 Perturbation paradigm. a*** | *We performed in-silico perturbations by in- and decreasing the individually estimated Peri Exc. for T1. For each perturbation, we calculated the resulting FC and FCDs and compared them to the patient's empirical FC and FCDs after one year (T3) by calculating the error between them. Whereas some patients required an increase (upper right) or a decrease (lower right) of Peri Exc. to approximate the brain dynamics and connectivity of T3, some patients did not show clear differences (lower left). As statistical benchmark we considered a narrow range around the initial Peri Exc. for T1 (blue box; see Methods) and compared it to an equivalently sized interval centred on the Peri Exc. value (red dot) that minimizes the error towards T3.*
